# Nucleotide metabolism is linked to cysteine availability

**DOI:** 10.1101/2022.09.30.510321

**Authors:** Annamarie E. Allen, Yudong Sun, Fangchao Wei, Michael A. Reid, Jason W. Locasale

## Abstract

The small molecule erastin inhibits the cystine-glutamate antiporter, system x_c_^-^, which leads to intracellular cysteine and glutathione depletion. This can cause ferroptosis, which is an oxidative cell death process characterized by uncontrolled lipid peroxidation. Erastin and other ferroptosis inducers have been shown to affect metabolism but the metabolic effects of these drugs have not been systematically studied. To this end, we investigated how erastin impacts global metabolism in cultured cells and compared this metabolic profile to that caused by the ferroptosis inducer RSL3 or in-vivo cysteine deprivation. Common among the metabolic profiles were alterations in nucleotide and central carbon metabolism. Supplementing nucleosides to cysteine-deprived cells rescued cell proliferation in certain contexts, showing that these alterations to nucleotide metabolism can affect cellular fitness. While GPX4 inhibition caused a similar metabolic profile as cysteine deprivation, nucleoside treatment did not rescue cell viability or proliferation under RSL3 treatment, suggesting that these metabolic changes have varying importance in different scenarios of ferroptosis. Together, our study shows how global metabolism is affected during ferroptosis, and points to nucleotide metabolism as an important target of cysteine deprivation.

## Introduction

Cysteine is a sulfur-containing amino acid which is essential for glutathione biosynthesis and redox homeostasis^1–3^. It can be synthesized from methionine through the transsulfuration pathway^4^ or imported as its oxidized form cystine via the system x_c_^-^ amino acid antiporter which exchanges cystine for glutamate^5^. However, many cancer cells cannot meet their requirements for cysteine through the transsulfuration pathway^6^, and certain cancers have been shown increased reliance on the system x_c_^-^ transporter^7–10^.

Inhibition of system x_c_^-^ can lead to ferroptosis, which was described over ten years ago as an oxidative, iron-dependent form of cell death that was induced by the small molecule erastin^11^. Erastin inhibits system xc- and causes cysteine and glutathione depletion^11–13^. Because glutathione is a cofactor for the glutathione peroxidase enzyme GPX4, decreased levels of glutathione can diminish the ability of GPX4 to detoxify potentially destructive lipid peroxide species^14^. Ferroptosis is defined as the cell death that occurs when cells are unable to adequately detoxify these species, whether that be due to glutathione depletion or direct loss of GPX4 function.

Ferroptosis contributes to pathological cell death in numerous disease models including those of glutamate-induced neurotoxicity, Huntington’s disease, periventricular leukomalacia, renal tubular injury and acute renal failure, ischemia/reperfusion injury, and liver damage^11,15–21^. Additionally, ferroptosis has shown considerable significance in the context of cancer^22–24^. This is partially due to studies showing many drug-resistant cell types are sensitive to erastin and the GPX4 inhibitor RSL3, or that erastin can enhance the effectiveness of chemotherapy and radiotherapy ^12,22,25–30^. There are also data which suggests certain cancers have increased reliance on system x_c_^-7–9^, prompting the investigation of whether its blockade could be a selective cancer therapy^31^.

Altered metabolism particularly from the mitochondria has been shown to influence ferroptosis providing additional aspects of the process^32^, but whether this picture is complete is unknown. To investigate this question, we used our metabolomics platform, which offers a broad coverage of cellular metabolism by measuring over 300 metabolites from more than 40 different metabolic pathways^33,34^ and found that changes in nucleotide metabolism accounted for many of the largest ferroptosis-induced alterations observed. Alterations in central carbon metabolism, including glycolysis and the citric acid cycle were also observed, and suggest that defective energy production may lead to altered nucleotide levels.

## Results

### Alterations in nucleotide metabolism are overrepresented in metabolic profile of ferroptosis

To determine how global metabolism is affected when cells are undergoing system x_c_^-^ or GPX4 inhibition, we treated three different ferroptosis-sensitive cell lines with an IC_50_ dose of ferroptosis inducers erastin or RSL3 (Fig. 1A) for 15 hours (before the onset of cell death) and then extracted polar metabolites for LC-MS based metabolomics and metabolic profiling. As expected, alterations in cysteine-related metabolites were observed in the metabolic profile of erastin-treated cells (Fig. 1B, D, F). However, all three cell lines also showed many alterations in the levels of nucleotide-related metabolites including nucleotides, nucleotide precursors and breakdown products, and/or nucleotide-sugars in both erastin and RSL3-treated cells, and these were some of the largest metabolic alterations observed (Fig. 1 B-G). To determine whether nucleotide-related metabolites were statistically overrepresented in the metabolic profile of erastin and RSL3-treated cells, metabolites with p<0.05 on each volcano plot were sorted by the absolute value of the difference between DMSO and drug-treated. The largest 10% of these metabolites were classified as nucleotide-related or not, and the percentage that were nucleotide related is shown in Fig. 1 H, I. Statistical overrepresentation of nucleotide-related metabolites was determined by comparing this percentage to the percentage of nucleotide-related metabolites that would be expected in the top 10% based on chance. Nucleotide-related metabolites were statistically overrepresented in erastin-treated BT-549 and HT-1080 cells, and in RSL3-treated HT-1080 and U2OS cells (Fig. 1 H, I). In erastin-treated U2OS and RSL3-treated BT-549 cells, nucleotide-related metabolites were not statistically overrepresented, but nucleotide changes were still observed in the metabolic profile (Fig. 1 E, F). Common nucleotide alterations observed included increased nucleoside levels (Fig. S1 A-C), decreased nucleotide precursor levels (Fig. S1 D-F) and decreased nucleotide di- and triphosphate levels (Fig. S1 G-O). To determine whether these alterations could be the result of off-target effects of either drug, we co-treated HT-1080 cells with erastin or RSL3 and ferroptosis inhibitors deferoxamine (DFO), ferrostatin-1 (ferrostatin), or the lipophilic antioxidant Trolox. We found that nearly all nucleotide-related metabolic alterations were at least partially rescued by ferrostatin treatment (Fig. S2 A), and most of the alterations were also partially rescued by DFO or Trolox treatment. We also found that ferrostatin fully rescued the majority of RSL3-induced nucleotide alterations, and fully rescued 50% of erastin-induced nucleotide changes, while DFO fully rescued 42 and 46% of RSL3 and erastin-induced nucleotide changes, respectively (Fig. S2 B). DFO was notably less effective than ferrostatin at rescuing nucleotide alterations, however this is unsurprising as DFO has been shown to affect nucleotide metabolism itself^35^. The lesser effectiveness of Trolox compared to ferrostatin was also not surprising, as ferrostatin has been shown to more efficiently suppress lipid peroxidation in the context of ferroptosis than Trolox^36^. These data show that many of the observed nucleotide alterations were reversed by validated ferroptosis inhibitors and are unlikely the result of off-target effects of either drug.

**Figure 1.**
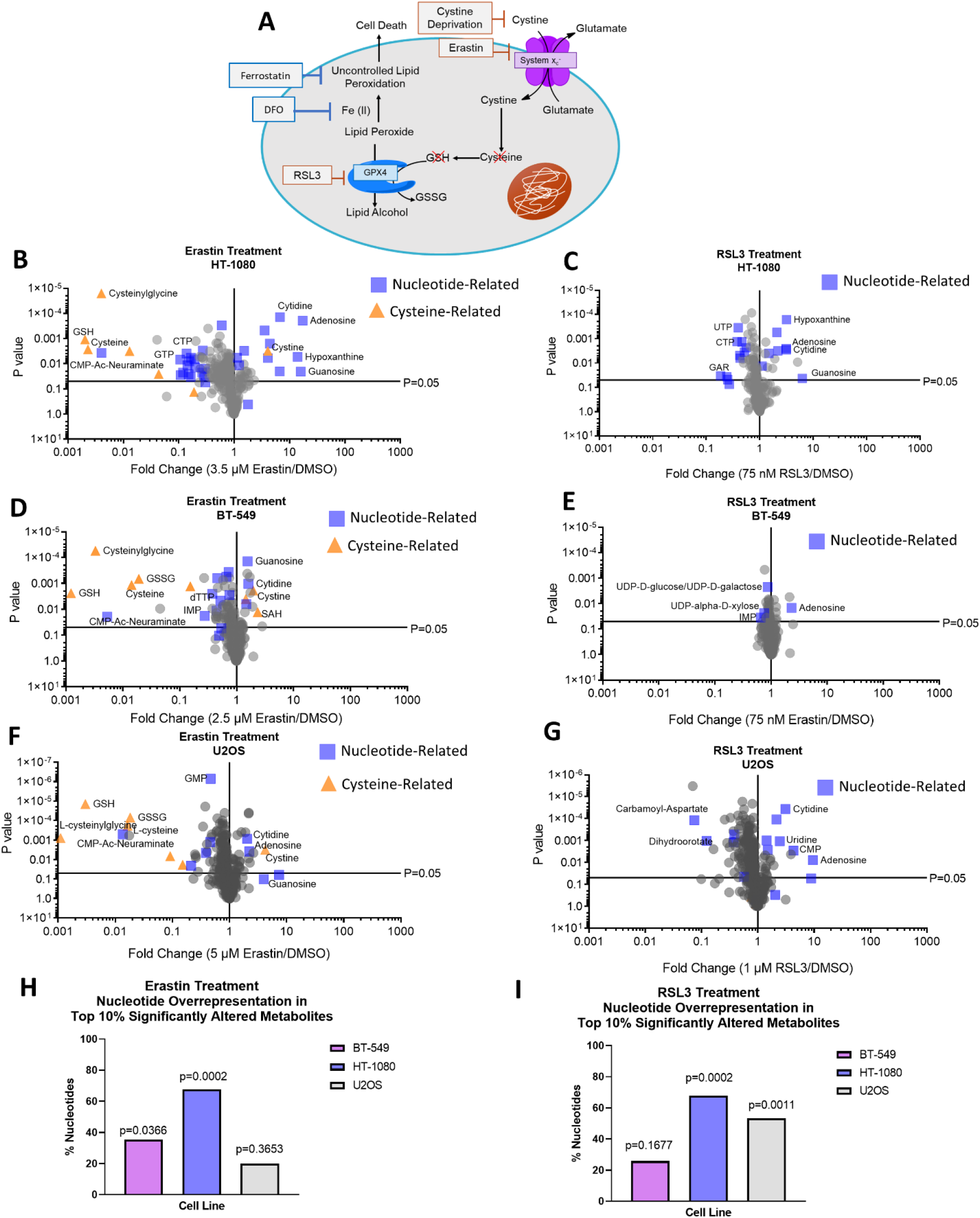
Alterations in nucleotide metabolism are overrepresented in metabolic profile of ferroptosis. A) Schematic of ferroptosis pathway B-I) Results from polar metabolomics shown as volcano plots using p-values generated from multiple unpaired t-tests on log-transformed ion intensity values. H, I) The percentage of the largest 10% of significantly altered metabolites that were nucleotides, nucleotide precursors and breakdown products, and nucleotide-sugars/lipids. For each volcano plot, metabolites with p<0.05 were sorted by the absolute value of the difference between DMSO and drug treatment, and the largest 10% were categorized as nucleotide-related or not. The percentage of these that were classified as nucleotide-related is graphed and the p-value is from a one-sided Fisher’s exact test comparison of this percentage versus the expected number of nucleotide-related metabolites that would be in the top 10% most significantly altered based on chance. Abbreviations: Deferoxamine (DFO), Reduced glutathione (GSH), Oxidized glutathione (GSSG), Glycineamide ribonucleotide (GAR), S-Adenosyl-L-homocysteine (SAH), CMP-N-Acetyl-Beta-Neuraminate (CMP-Ac-Neuraminate)

### Erastin decreases nucleotide biosynthesis

Decreased nucleotide levels could result from insufficient nucleotide biosynthesis, insufficient energy levels, or both. To better understand how the ferroptosis inducer erastin causes this pattern of nucleotide changes, we performed ^13^C-glucose tracing in BT-549 and HT-1080 cells treated with erastin and examined how nucleotide biosynthesis and energy producing pathways were affected. Universally labeled ^13^C glucose (U-^13^C glucose) labels purines and pyrimidines through conversion to ribose-5-phosphate (R5P) via the pentose phosphate pathway (Fig. 2A). M+5 R5P showed no differences in fractional labeling between DMSO- and erastin-treated cells (Fig. 2 B, C), suggesting that pentose phosphate pathway defects are not responsible for any decreased m+5 nucleotide labeling. In HT-1080 cells, purines did not show decreased m+5 fractional labeling at the timepoint tested (Fig. 2 D-F), but pyrimidines did (Fig. 2 G-J). In BT-549 cells both purines and pyrimidines showed decreased m+5 fractional labeling (Fig. 2K-O). These results show that erastin disrupts nucleotide biosynthesis, which may partially explain why some nucleotides are depleted in cells treated with erastin.

**Figure 2.**
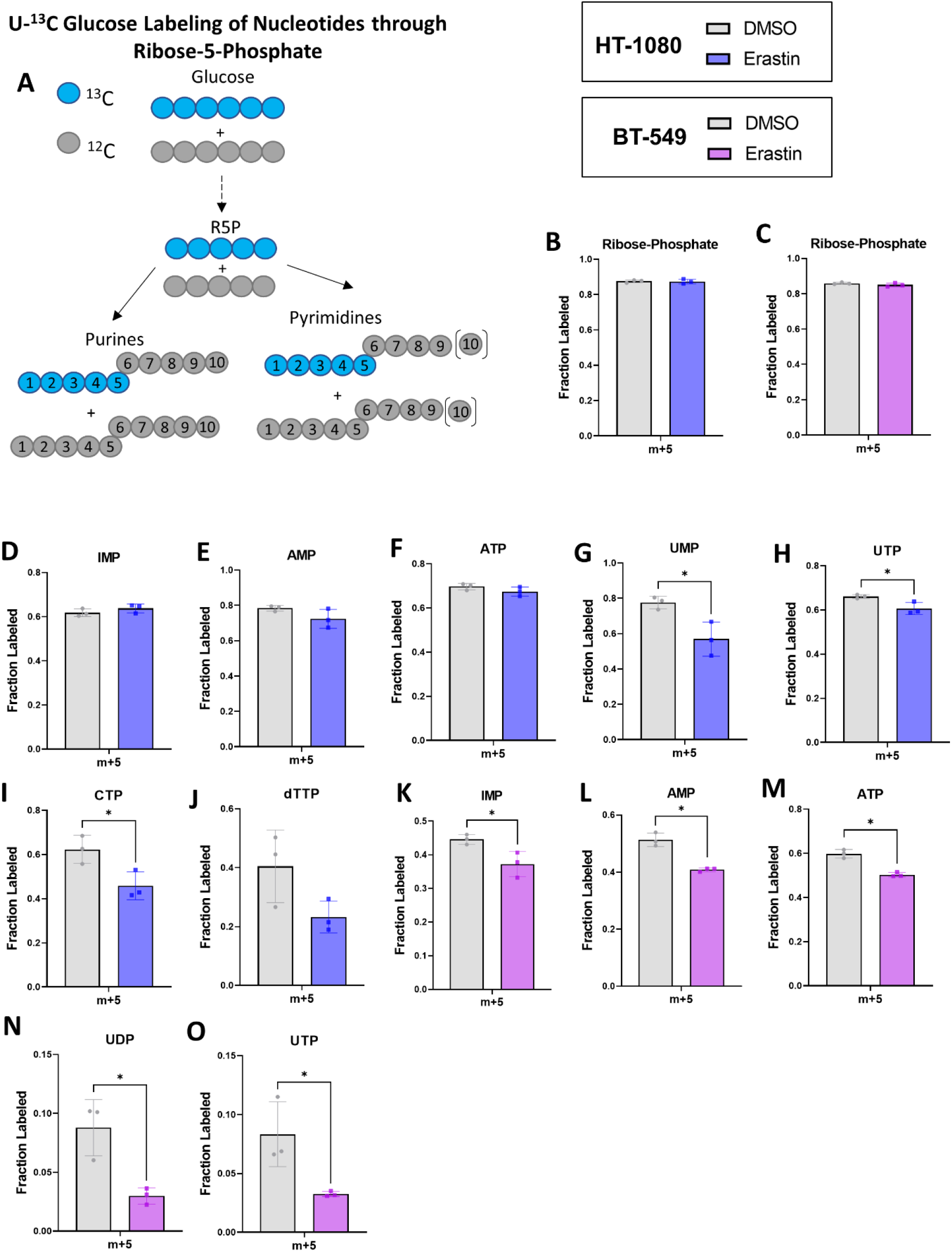
Erastin decreases nucleotide biosynthesis. A) Schematic showing U-13C Glucose nucleotide labeling through m+ 5 ribose-5-phosphate. B, D-H). M+5 fractional labeling plotted for relevant nucleotides and precursors from HT-1080 cells treated with 2.5 uM erastin. C, K-O) M+5 fractional labeling plotted for relevant nucleotides and precursors collected from BT-549 cells treated with 3.5 uM erastin. For Figs. B-O, an unpaired T-test was performed to compare DMSO to erastin for each metabolite. * indicates p<0.05 for this test, no stars shown indicates p>0.05.

### Erastin alters glycolysis and citric acid cycle activity and decreases energy levels

Insufficient energy levels may also explain why higher-energy nucleotide di- and triphosphates were the type of nucleotide most often depleted under erastin treatment (Fig. S1 J-O), while lower-energy nucleotide monophosphate (Fig. S1 G-I) and nucleoside (Fig. S1 A-C) levels either increased or showed smaller fold-changes than the di- and triphosphates. To determine whether this could be occurring, we first looked at the total levels of metabolites in the energy-producing pathways glycolysis and the citric acid cycle (TCA). Erastin-treated HT-1080 and BT-549 cells showed significant depletions in many glycolytic intermediates (Fig. 3 A, D), and in TCA intermediates (Fig. 3 B, E). The ratio of ATP/AMP, an indicator of cellular energy status, was also significantly decreased in both cell lines (Fig. 3 C, F). The NADH/NAD+ ratio, another indicator of energy status, was also decreased, although not significantly (Fig. S3 B, F), and increased protein levels of phosphorylated AMP-activated protein kinase (pAMPK) were observed (Fig. S3J), further suggesting there is an energy depletion under erastin treatment. U-^13^C glucose tracing (Fig. S3A) showed that pyruvate and lactate had significantly decreased m+3 fractional labeling under erastin treatment in both cell lines (Fig. 3 G, H, K, L) and a decreased m+3 pool size (Fig. S3 C, D, G, H), which indicates less glycolytic activity over time.

**Figure 3.**
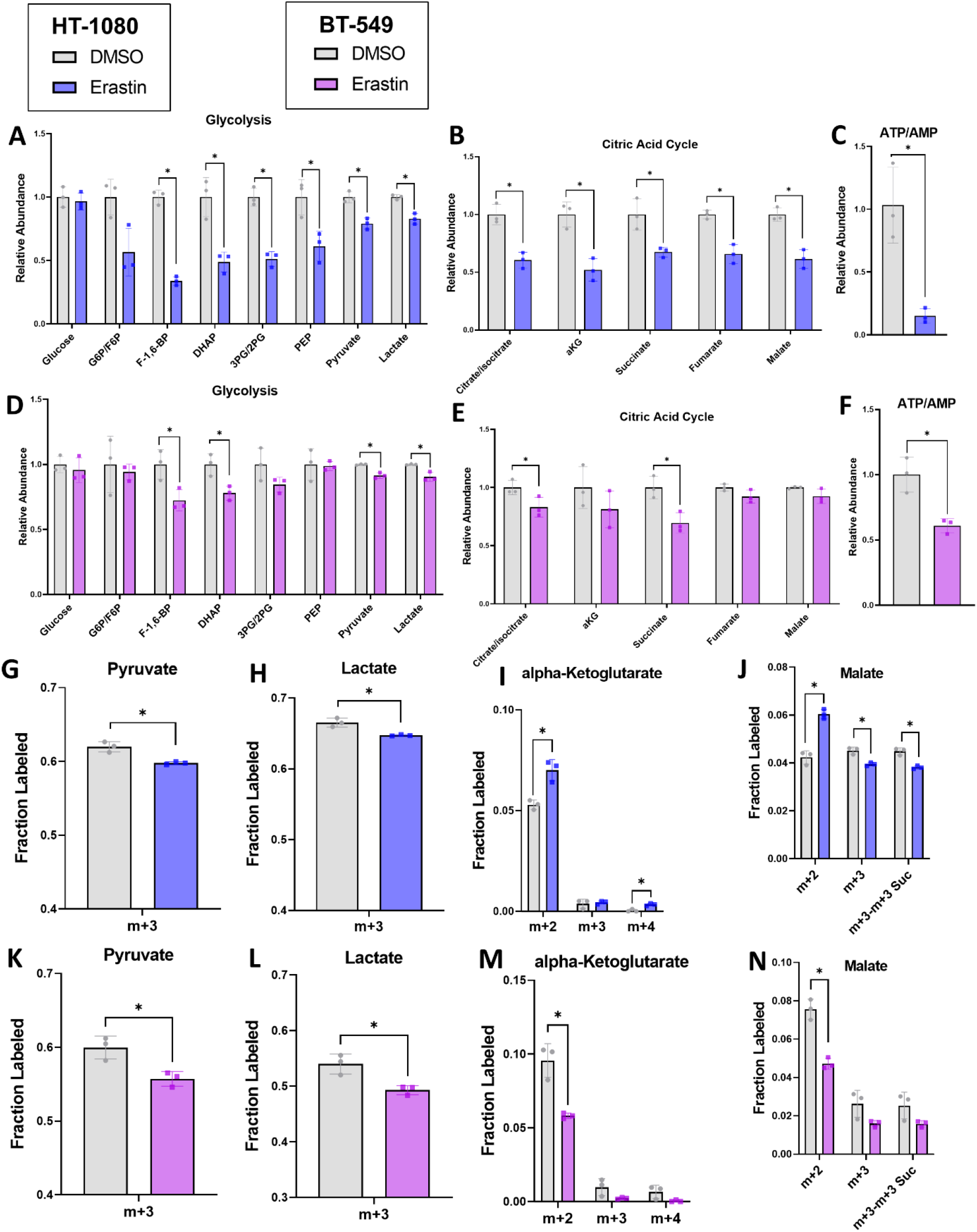
Erastin alters glycolysis and citric acid cycle activity and decreases energy levels. A-F) Relative levels of metabolites from relevant pathways in HT-1080 cells treated with 2.5 μM erastin or BT-549 cells treated with 3.5 μM erastin. G-N) Fractional labeling plotted for relevant isotopomers in glycolysis and TCA in HT-1080 cells treated with 2.5 μM erastin or BT-549 cells treated with 3.5 μM erastin and subjected to U-^13^C Glucose tracing. In Figs. J, N m+3-m+3 Suc represents the m+3 fraction of malate minus the m+3 fraction of succinate, which represents glucose entry into the TCA via pyruvate carboxylase. For Figs. A-B and D-E a two-way repeated measures ANOVA was performed on log-transformed values and followed by an uncorrected Fisher’s LSD test if interaction term or main effect of treatment p<0.05. * indicates p<0.05 when comparing DMSO to Erastin treatment in Fisher’s LSD test, no stars shown indicates p>0.05. For Figs. C, F, G-N an unpaired T-test was performed to compare DMSO to erastin treatment. If more than one isotopomer is shown on the graph, a separate uncorrected T-test was performed for each. * indicates p<0.05 for this test, no stars shown indicates p>0.05.

TCA cycle labeling showed a different pattern in the two different cell lines, with significantly increased ^13^C glucose-derived m+2 labeling on the TCA cycle intermediates alpha-ketoglutarate (aKG) and malate in erastin-treated HT-1080 cells (Fig. 3 I, J) and significantly less in erastin-treated BT-549 cells (Fig. M, N). M+2 aspartate, which is produced from the TCA cycle, also showed increased m+2 fractional labeling in erastin-treated HT-1080 cells and decreased m+2 fractional labeling in erastin-treated BT-549 cells (Fig. S3 E, I). M+4 aKG, which is produced in the second turn of the TCA cycle (Fig. S3A), was also significantly increased in erastin-treated HT-1080 cells (Fig. 3I) and non-significantly decreased in erastin-treated BT-549 cells (Fig. 3M). Before entering the TCA cycle, glucose is converted to pyruvate, which can enter the TCA via two enzymes, pyruvate dehydrogenase (PDH) and pyruvate carboxylase (PC) (Fig. S3A). Pyruvate entry into the TCA via PC serves an anaplerotic role and can be distinguished from PDH-derived pyruvate labeling by subtracting the m+3 fraction of succinate from the m+3 fraction of malate or aspartate^37,38^. Interestingly, the m+3 malate fraction minus the m+3 succinate fraction was decreased in both cell lines treated with erastin, although only significantly in HT-1080 cells (Fig. 3J, N). The m+3 aspartate minus the m+3 succinate fraction was not decreased in erastin-treated HT-1080 cells (Fig. S3E), but the labeling pattern of malate is more reliable to interpret since aspartate labeling is confounded by aspartate in the culture media^38^. Together, these results show that erastin-treated HT-1080 cells increase glucose usage into the TCA via PDH, while decreasing anaplerotic glucose entry into the TCA via PC. Erastin-treated BT-549 cells show decreased glucose usage in the TCA cycle through both PDH and PC indicating mitochondrial plasticity.

### Exogenous nucleosides rescue diminished cell proliferation under erastin treatment

To understand whether depleted nucleotide levels have a functional significance in erastin-induced cell death and/or growth reduction, we co-treated cells with erastin and a nucleoside cocktail (Fig. 4A) that has previously been used to increase viable cell number in cells with diminished nucleotide production^39^. We found that co-treating with exogenous nucleosides caused a small increase in relative viable cell number in erastin-treated HT-1080 cells (Fig. 4B), but a larger nucleoside rescue effect was seen in HT-1080 cells when directly depleting cystine in the media (Fig. 4C). Nucleosides also partially rescued relative viable cell number and restored proliferation in erastin-treated BT-549 cells (Fig. 4D-F). Because nucleosides increased the number of viable cells 3 days after treatment in erastin-treated cells, but not 1 day after treatment (Fig. 4E) the nucleosides likely increase viable cell number by restoring proliferation rather than by inhibiting cell death in erastin-treated cells. Nucleosides did not show any rescue effect in in erastin-treated U2OS cells (Fig. S4 G) or in RLS3-treated HT-1080 or BT-549 cells (Fig. S4 H, I), while ferrostatin co-treatment caused a large rescue effect in every cell line tested that was treated with erastin and RSL3 (Fig. S4 A-F). This suggests that the importance of nucleotide-related changes in affecting cellular fitness during ferroptosis is likely secondary to other factors and plays a role in certain circumstances. To understand how nucleosides increase cell proliferation in these circumstances, we measured cysteine, glutathione, and nucleotide levels in BT-549 cells treated with erastin with or without nucleoside co-treatment for 15 hours and found that nucleosides do not alter the level of cysteine or glutathione depletion that is caused by erastin (Fig. 4G). Nucleoside co-treatment did significantly increase the levels of certain nucleotides, including GMP, CMP, (Fig. 4 H) CDP, (Fig. 4I) and CTP (Fig. 4J), while others including ADP, UDP, ATP, and dTTP were unaffected. While the levels of several nucleosides tended to increase when cells are treated with ferroptosis inducers alone (Supplementary Figure 1A-C), this amount of increase is small in comparison to when exogenous nucleosides are supplemented into the media (Supplementary Figure 4J), suggesting that the higher nucleoside levels in the erastin only condition are not enough to provide sufficient nucleotide salvage material. These results show that exogenous nucleosides can partially rescue nucleotide levels and cell proliferation which are decreased by erastin in certain cell lines and conditions, suggesting that nucleotide depletion plays a role in the loss of cellular fitness caused by erastin.

**Figure 4.**
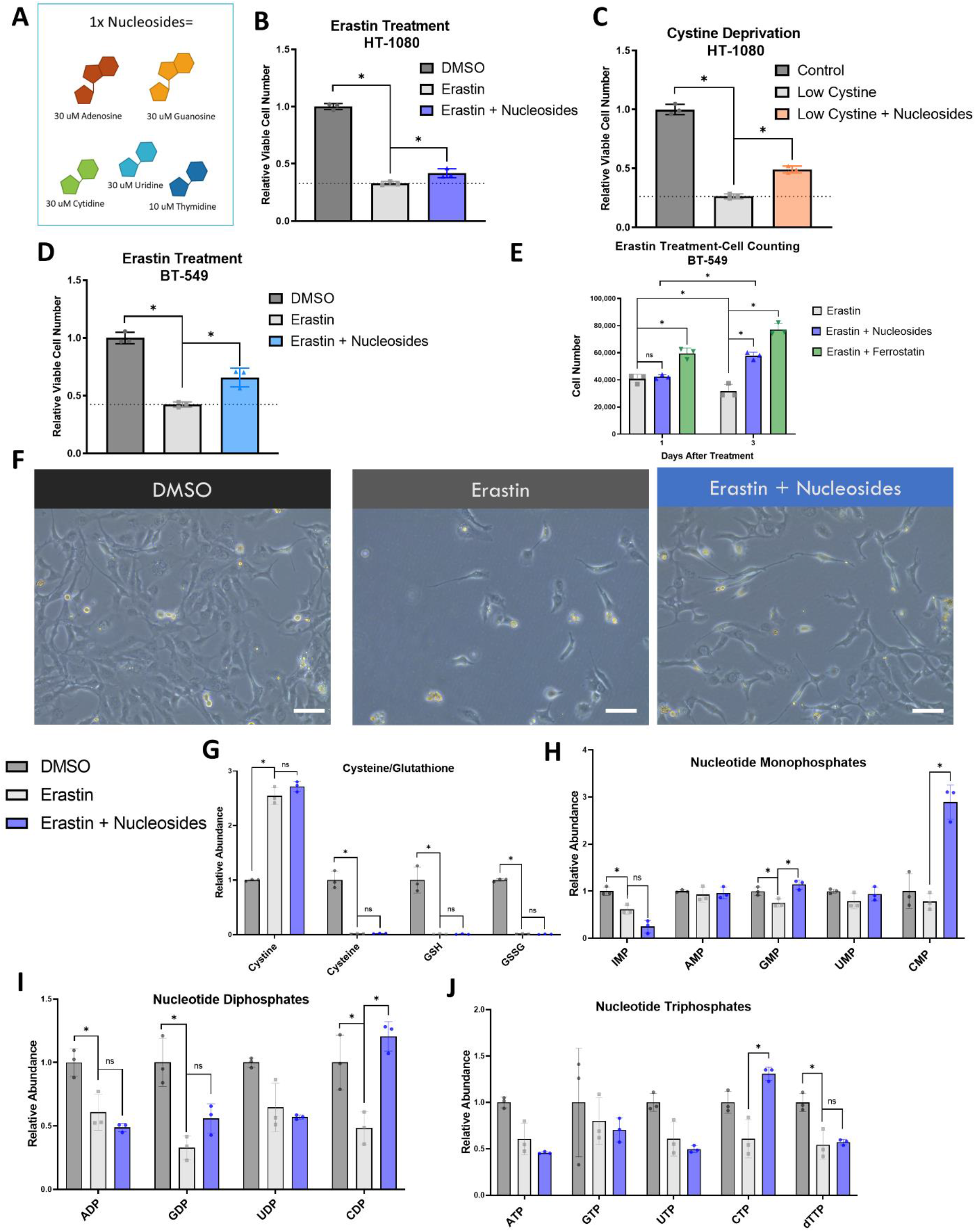
Exogenous nucleosides rescue diminished cell proliferation under erastin treatment. A) Identity and concentration of nucleosides in “1x Nucleosides” cocktail. B) Relative viable cell number in HT-1080 cells treated +/- 2.5 μM erastin and 1x nucleosides for 48 hours. C) Relative viable cell number in HT-1080 cells cultured in control media or media containing low cystine +/- 1x nucleosides for 48 hours. D) Relative viable cell number in BT-549 cells treated +/- 3.5 μM erastin and 1x nucleosides for 48 hours. E) Cell number as determined by an automated cell counter 24 and 72 hours after treatment with 3.5 μM erastin +/- 1x nucleosides and 0.5 μM Ferrostatin. F) Pictures of BT-549 cells taken after 48 hours +/- 3.5 μM erastin and 1x nucleosides. The scale bar shown is equal to 100 μm G-J) Relative levels of selected metabolites from BT-549 cells treated with 3.5 μM erastin +/- nucleosides. For Figs. B-D, a one-way ANOVA was performed and followed by an uncorrected Fisher’s LSD test when p<0.05 for ANOVA. * indicates p<0.05 in Fisher’s LSD test. For E, a two-way ANOVA was performed and followed by an uncorrected Fisher’s LSD comparing erastin to erastin + nucleosides and erastin + ferrostatin at each timepoint. Erastin and Erastin + Nucleosides at Day 1 was also compared to Day 3. For Figs. H-K, a two-way repeated measures ANOVA was performed on log-transformed values and followed by an uncorrected Fisher’s LSD test if interaction term or main effect of treatment p<0.05. * indicates p<0.05 when comparing Erastin to DMSO or to Erastin + Nucleosides. Bar is plotted at mean and error bars show standard deviation. Each dot represents one biological replicate.

### Nucleotide and central carbon metabolism are altered in flies fed a cysteine-deprived diet

We next sought to explore an in-vivo model of cysteine and glutathione depletion, which could be relevant to ferroptosis, to determine: 1) whether glutathione depletion can be achieved through dietary means and 2) whether similar nucleotide alterations are observed *in vivo* when cysteine and glutathione are depleted. We speculated that long-term cysteine and glutathione depletion would decrease fly lifespan. Since cysteine is a conditionally essential amino acid and can be synthesized from methionine, we did not know whether dietary cysteine depletion alone would affect fly survival. We therefore depleted cysteine, methionine, or both in the food of male or virgin female *w^1118^ Drosophila melanogaster* and assessed survival as compared to flies fed a control chemically defined diet. In male flies, cysteine depletion alone did not affect survival and methionine depletion alone slightly increased median survival, but the depletion of both dietary cysteine and methionine significantly decreased median survival (Fig. 5A and B). In female flies, we saw similar patterns, but changes were not statistically significant (Fig. S5A, B), so we decided to investigate potential dietary-induced metabolic changes in male flies. We fed male flies a control or cysteine and methionine free chemically defined diet for three weeks, and then collected fly bodies and heads for separate metabolic profiling. Fly bodies and heads from flies fed the cysteine and methionine-free diets showed significantly depleted cysteine and glutathione levels (Fig. 5C and S5C), demonstrating that diet-mediated glutathione depletion is possible. Next, we assessed whether nucleotide alterations occurred in cysteine and methionine-deprived flies and found significant depletions in nucleotide monophosphates in both the fly head (Fig. S5E) and body (Fig. 5E). We also found depletions in nucleotide diphosphates that were significant in the fly body (Fig. 5F) but not the head (Fig. S5F). ATP was the only nucleotide triphosphate that was measured in the flies, and it was depleted in both the head and body, but not to a statistically significant level (Fig. S5G and 5G). We also observed significant differences in citric acid cycle intermediates in both the head and body (Fig. 5H and S5H) and significantly depleted NADH levels in the body (Fig. 5I). We wondered whether supplementing nucleosides into the diet of cysteine and methionine-deprived flies could alter survival. We did not know whether this strategy would be effective *in vivo* since it would depend on the digestion, absorption, and distribution of dietary nucleosides throughout the body, and because nucleotide depletions may only play a small role in the decreased survival observed in cysteine and methionine-deprived flies. However, in light of these complications, supplementing 2.5 mM nucleosides into the diet caused a small but significant increase in survival of cysteine and methionine-deprived male flies (Fig. 5J). Together, these data show that long-term dietary cysteine and glutathione depletion decrease survival and trigger similar *in vivo* metabolic alterations involving nucleotide metabolism and the citric acid cycle as observed in cultured cells.

**Figure 5.**
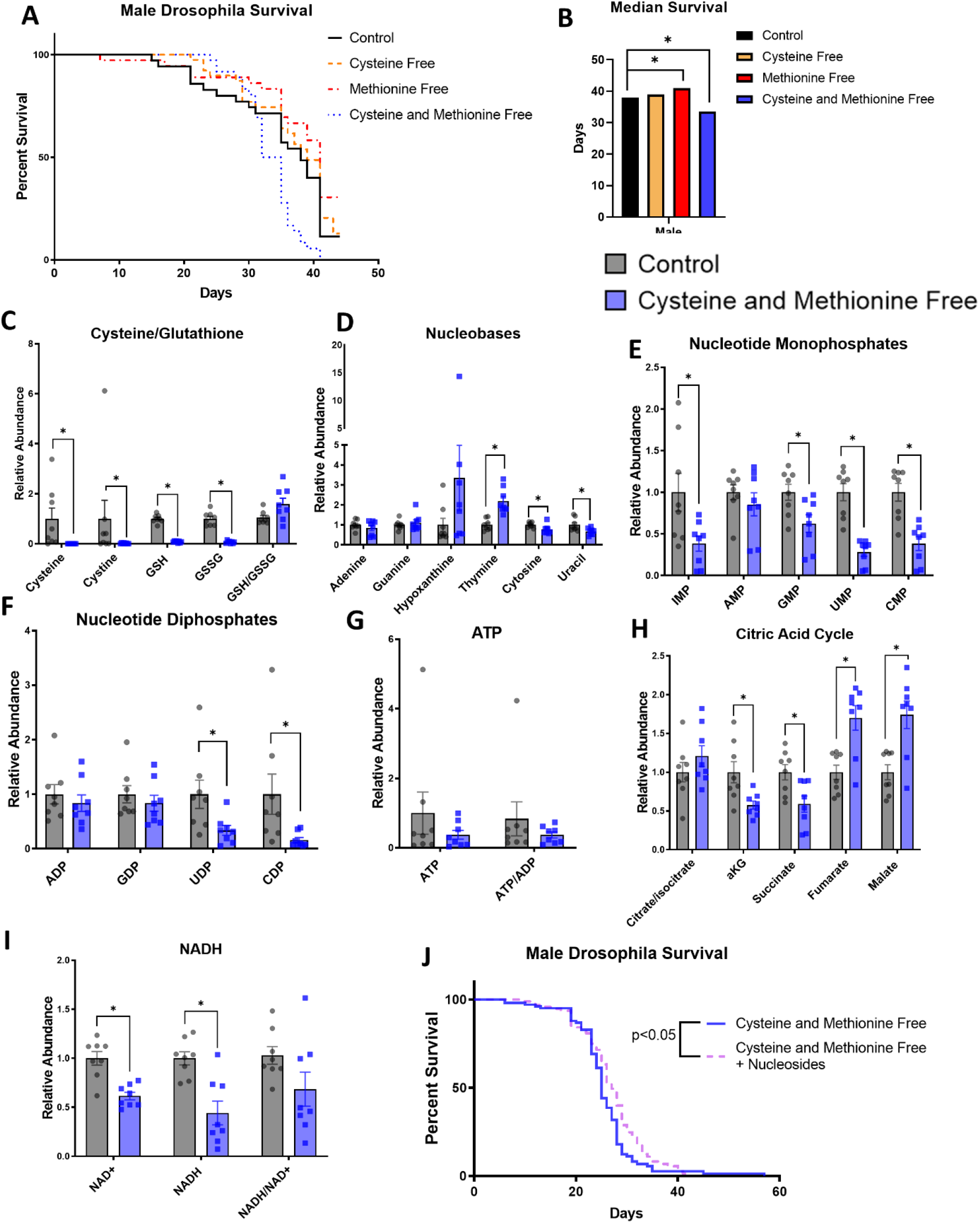
Nucleotide and central carbon metabolism are altered in male w1118 *Drosophila melanogaster* fed a cysteine-deprived diet. A) Male *w1118 drosophila* survival when fed chemically defined diets with or without cysteine and methionine. B) Median survival of male flies from survival curves shown in Fig. 4A.C-I) The relative abundance of selected metabolites from male *drosophila* bodies fed the indicated diets for 3 weeks. Each dot represents one fly. J) Male *w1118 drosophila* survival when fed chemically defined diets without cysteine and methionine +/- nucleosides. Nucleosides added into the diet were Inosine, Uridine, Adenosine, Guanosine, Cytidine, and Thymidine (2.5 mM). For Fig. B, multiple log-rank Mantel-Cox tests were performed to compare male fly survival differences on control versus cysteine/methionine free diets. * indicates p<0.05 for these tests, while no star indicates p>0.05. For Figs. C-I, a two-way repeated measures ANOVA performed on log-transformed values and followed by an uncorrected Fisher’s LSD test if interaction term <0.05. * indicates p<0.05 for Fisher’s LSD test. For Fig. J, a log-rank Mantel-Cox test was performed to compare fly survival on cysteine and methionine free versus cysteine and methionine free + nucleosides diets.

## Discussion

Previous studies have linked ferroptosis to nucleotide metabolism through dihydroorotate dehydrogenase (DHODH)^40^ and through ribonucleotide reductase (RNR)^41^. Here, we show that alterations in nucleotide metabolism account for some of the largest changes in the metabolome of cells treated with ferroptosis inducers erastin and RSL3. This metabolic profile included alterations which have previously been observed in RSL3-treated cells including depleted pyrimidine precursor N-carbamoyl-L-aspartate and elevated uridine^40^, but also showed a much wider set of nucleotide alterations in both erastin and RSL3-treated cells including elevated levels of other nucleosides such as adenosine, guanosine, and cytidine, and broadly depleted nucleotide di- and triphosphate levels.

We investigated two mechanisms through which the nucleotide alterations observed in erastin-treated cells could occur. Tracing with ^13^C glucose showed that erastin decreases m+5 labeling on pyrimidines in HT-1080 and BT-549 cells, and on purines in BT-549 cells. Because m+5 ribose was unaffected, decreased nucleotide biosynthesis under erastin treatment is not likely due to altered pentose phosphate pathway activity, and because nucleotides upstream of RNR showed altered labeling, RNR is also unlikely to cause altered nucleotide biosynthesis during erastin treatment. Decreased nucleotide biosynthesis can partially explain why nucleotide depletions occur during erastin treatment, but it does not explain why higher-energy nucleotide di- and triphosphates were depleted while lower-energy nucleotide monophosphates and nucleosides either accumulated or showed smaller depletions. It also does not explain why purines such ADP, GDP, ATP, and GTP were depleted in erastin-treated HT-1080 cells when purine biosynthesis was unaffected. Decreased levels of the ATP/AMP and NADH/NAD+ ratio in erastin-treated cells and induction of AMPK phosphorylation can better explain why these nucleotides are depleted. Decreased energy production can also affect nucleotide production because nucleotide biosynthesis requires significant energy input^42^.

Our study did not directly measure the source of energy depletion during erastin treatment, but the altered ^13^C glucose labeling seen on glycolysis and TCA intermediates shows that these energy-producing pathways were affected by erastin. Both cell lines showed decreased glycolysis over time under erastin treatment as indicated by altered pyruvate and lactate m+3 pools. This likely contributed to at least some of the energy depletion caused by erastin. Many metabolic enzymes, including several glycolytic and TCA enzymes, are affected by reactive oxygen species^43^, and the significant glutathione depletion caused by erastin treatment is likely to affect one or more of these enzymes through loss of redox homeostasis^44^. Lipid peroxidation has been observed in mitochondria^45,46^ and could also target enzymes in these and other pathways. A limitation of the study was only focusing on possible mechanisms of erastin-induced nucleotide alterations and not those of other ferroptosis inducers such as RSL3, or of genetic methods of ferroptosis induction.

Previous studies have found that increased ROS generation by mitochondria can contribute to ferroptosis^47,48^ and that erastin-induced ferroptosis can be inhibited by suppressing the TCA^45^. It has also been suggested that erastin increases mitochondrial metabolism through the opening of voltage-dependent anion channels in the outer mitochondrial membrane^47^. Our study found that TCA intermediates were depleted in erastin-treated HT-1080 and BT-549 cells. However, the two cell lines had opposite patterns in terms of ^13^C glucose labeling of TCA intermediates through pyruvate dehydrogenase, with HT-1080 cells showing increased glucose usage in the TCA and BT-549 cells showing decreased. Increased glucose usage in the TCA in erastin-treated HT-1080 cells may partially compensate for loss of glycolytic activity or decreased use of other TCA fuel sources. We did not determine whether blocking mitochondrial metabolism contributed to or suppressed erastin-induced cell death, but our results do show that mitochondrial metabolism is affected differently in different cell lines, and caution should be used when generalizing the results obtained from a single model.

Our results show that a major effect of erastin-treated cells is nucleotide depletion, and that supplementing cysteine-deprived cells with exogenous nucleosides can partially rescue diminished cell proliferation at intermediate doses of erastin. These results indicate that the nucleotide changes observed in the metabolic profile of erastin-treated cells have a functional impact on cellular fitness. Restored nucleotide levels likely improve the rates of macromolecule biosynthesis and repair^42^. Glutathione depletion by erastin and other methods has shown to synergize with numerous chemotherapies including those that interfere with nucleotide synthesis^49^. Our study offers a plausible mechanism for why these synergisms occur. RSL3-treated cells showed a very similar metabolic profile to erastin-treated cells, but exogenous nucleosides did not rescue viable cell number under RSL3 treatment in any cell line tested. We did not investigate why this is but speculate that it may have to do with differences in the onset, strength, and location of lipid peroxidation caused by erastin versus RSL3. The efficacy of whether the exogenous nucleoside cocktail rescues viable cell number and nucleotide levels in erastin-treated cells likely depends on several factors including the extent to which nucleotide depletions are caused by insufficient nucleotide biosynthesis and salvage versus insufficient energy levels, the speed and ability of cells to salvage the nucleosides from the cocktail, and the role reduced nucleotide levels play in decreasing cellular fitness in different cell lines and conditions.

Our study also showed that glutathione depletion can be achieved through dietary methods.

Long-term cysteine and methionine restriction significantly decreased the lifespan of male flies and had a similar effect on virgin female flies that was not statistically significant. We also observed similar nucleotide and TCA cycle changes in the metabolic profile of male flies subjected to a cysteine and methionine-free diet as seen in cultured cancer cells treated with erastin. Because ferroptosis has been implicated in neurological disease^19,50^, we questioned whether the metabolic profile of fly heads would differ from fly bodies but found almost identical metabolic changes. Supplementing nucleosides into the diet of the cysteine-deprived cells caused a small but statistically significant increase in median survival, suggesting that the nucleotide depletions observed in the metabolic profile of cysteine-deprived cells may play a role in their decreased survival.

In summary, our study shows that nucleotide metabolism is linked to cysteine metabolism and erastin-induced ferroptosis which can play a role in diminished cell proliferation in certain contexts. Further, it also establishes that glutathione depletion causes alterations to nucleotide synthesis through several mechanisms but not entirely through a single step involving an enzyme such as RNR or DHODH. Given these links many questions in disease biology such as cancer and neuropathology are open for further investigation.

## Materials and Methods

### Cell Culture

BT-549, HT-1080, and U2OS cells were purchased from the Duke Cell Culture Facility and cultured in RPMI 1640 (Gibco) supplemented with 10% heat-inactivated fetal bovine serum (Sigma, F2442). Cells were cultured in a 37 °C, 5% CO2 atmosphere. All cell lines tested negative for mycoplasma contamination.

### Cell Viability Assays

Cells were seeded at a density of 2-4 x 10^3^ cells per well in 96-well plates with water in the outside wells and allowed to adhere overnight. Cells were treated the following day by replacing the seeding media with 100 μL treatment media containing drug treatments or nutrient restricted media and any rescue agents. Cells were incubated in treatment media for the indicated period of time and then MTS reagent ((3-(4,5-dimethylthiazol-2-yl)-5-(3-carboxymethoxyphenyl)-2-(4-sulfophenyl)-2H-tetrazolium) purchased from Abcam, (ab197010) was mixed 1:1 with sterile PBS and 20 μL of the mixture was added to each well. Cells were incubated with MTS reagent for 2 hours, and after brief shaking absorbance was read at 490 nm on a microplate reader.

### IC_50_ Dose Determinations

For each cell line, IC_50_ doses of erastin and RSL3 were determined by seeding cells at a density of 2-4 x 10^3^ cells per well in 96-well plates with water in the outside wells. Doses of erastin ranging from 0-50 μM were added to wells in triplicate, and doses of RSL3 ranging from 0-1.5 μM were added to wells in triplicate. Cells were incubated with erastin/RSL3 for 24 hours, and then MTS cell viability assays were performed as previously described. IC_50_ dose curves were plotted and an IC_50_ dose for each drug in each cell line was calculated based off the curve.

### Drugs and Rescue Agents

Erastin (17754), RSL3 (19288), and Ferrostatin-1 (17729) were purchased from Cayman Chemical. The “1x nucleosides” were EmbryoMax Nucleosides purchased from MilliporeSigma (ES-008-D).

### Cystine Deprivation

Low cystine media was made from RPMI without L-glutamine, L-cystine, L-methionine, and L-cysteine (MP Biomedicals 1646454) supplemented with 10% dialyzed heat-inactivated fetal bovine serum (VWR SH30079.03). L-methionine (Amresco E801) and L-glutamine (Amresco 0374) were added back to the media to the concentration found in RPMI 1640 and L-Cystine (Amresco J993) was added back to control media but not to cystine free media. Both control and cystine free media were adjusted to pH 7.4 and then sterilized using a 0.10 μm filter (MilliporeSigma S2VPU02RE). To find an equivalent level of cystine deprivation that matched the level of cell viability/proliferation reduction caused by erastin in HT-1080 cells, we added varying levels of control media to the cystine free media and measured cell viability after 24 hours (the same way IC_50_ doses of erastin were found) and selected 7.5% cystine for the low-cystine media used in the rescue experiment.

### Cell Counting

Cells were seeded at a density of 4×10^4^ cells per well in six-well plates and allowed to adhere overnight. Cells were treated the following day by replacing media with 2 mL treatment media containing drug treatments and rescue agents. After 1 or 3 days of treatment, cells were trypsinized and then counted on a MOXI Z automated cell counter (Orflo).

### Microscopy

Cells were seeded at a density of 8×10^4^ cells per well in six-well plates and allowed to adhere overnight prior to treatment. Cells were incubated in treatment media for 48 hours prior to imaging. Images were captured using a Leica DM IL LED microscope equipped with a Leica MC170HD camera at x10 objective using LAS EZ software (Leica). Scale bars = 100 μm.

### Western Blotting

Total cell protein was extracted with lysis buffer (RIPA, Sigma-Aldrich, catalog no. R0278) containing 1% protease-phosphatase inhibitor cocktail (Thermo Scientific, catalog no. PI78440). Protein concentrations were measured with the BCA Protein Assay (Thermo Fisher Scientific, catalog no.23224). Total protein was resolved on 10% SDS–polyacrylamide gel electrophoresis gels and transferred onto PVDF membranes (Bio-Rad, catalog no. 1704156) by Trans-Blot Turbo (Bio-rad). After 5% BSA blocking, the membranes were incubated with primary antibodies containing AMPKα (Cell Signaling Technology, catalog no. 2532), Phospho-AMPKα (Thr172) (Cell Signaling Technology, catalog no. 2531) and β-actin (Cell Signaling Technology, catalog no. 4970) overnight at 4°C separately. Secondary antibody (Cell Signaling Technology, catalog no. 7074) was probed for 1 hour at room temperature. The chemiluminescence signal intensity of protein was detected with chemiluminescence (Thermo Scientific, catalog no. PI34096) and the ChemiDoc Touch Imaging System (Bio-rad).

### Stable Isotope Labeling

Cells were seeded at 8 x 10^4^ cells per well into six-well plates and allowed to adhere overnight prior to treatment. Treatment media was added to cells for 4 hours, and then replaced with U-^13^C Glucose-containing treatment media, which cells were cultured in for 12 hours. The U-^13^C Glucose-containing treatment media was made from glucose-free RPMI (Gibco 11879-020) supplemented with 10% heat-inactivated fetal bovine serum (Sigma, F2442) with [U-13C] glucose (Cambridge Isotope Laboratories, #CLM-1396) added at 2 g/L. Metabolites were then extracted and measured via LC-MS according to the protocols below.

### Polar Metabolite Extraction from Cells

Polar metabolite extraction was conducted as previously described^33,34^. Briefly, 8-12 x 10^4^ cells per well were seeded into six-well plates and allowed to adhere overnight prior to treatment.

Cell confluence was equal across conditions at the time of extraction. Following treatment, medium was quickly aspirated, plates were placed on dry ice, and 1 mL of 80% methanol/water extraction solvent (Optima LC-MS grade, Fisher; methanol, #A456; water, #W6) pre-cooled to −80°C was immediately added to each well prior to transferring the plates to −80°C for 15 min. The plates were then removed, placed on dry ice, and the cells were scraped into the extraction solvent and transferred to Eppendorf tubes. Metabolite extracts were then centrifuged at 20,000 × g at 4 °C for 10 min. The solvent in each sample was then transferred to a new Eppendorf tube and evaporated using a speed vacuum.

### Polar Metabolite Extraction from Flies

Flies were immobilized on ice and fly heads were separated from the thorax and abdomen (“body”) using tweezers, then both the head and both were flash-frozen in Eppendorf tubes using liquid nitrogen. Two heads a diet group were pooled together to have enough material for extraction. During extraction, the Eppendorf tubes were placed on dry ice and 200 μL 80% methanol pre-cooled to −80°C was added to each tube. The fly bodies and pooled heads were then homogenized using a tissue homogenizer. After homogenization, an additional 300 μL pre-cooled 80% methanol was added to each tube and tubes were vortexed and then centrifuged at 20,000 × g at 4 °C for 10 min. The solvent in each sample was then transferred to a new Eppendorf tube and evaporated using a speed vacuum. (note: solvents were the same as used in Polar Metabolite Extraction from Cells.

### Liquid chromatography

For polar metabolite analysis, the evaporated cell extracts were first dissolved in 15 μL LC-MS grade water and then15 μL methanol/acetonitrile (1:1 v/v) (Optima LC-MS grade, Fisher; methanol, #A456; acetonitrile, #A955) was added. Finally, samples were centrifuged at 20,000 × g at 4 °C for 10 min, and the supernatants were transferred to LC vials prior to HPLC injection (3 μL).

An XBridge amide column (100 x 2.1-mm inner diameter, 3.5 μm; Waters) was used on a Dionex (Ultimate 3000 UHPLC) for compound separation at room temperature. Mobile phase A was water with 5 mM ammonium acetate, pH 6.9, and mobile phase B was 100% acetonitrile. The gradient is linear as follows: 0 min, 85% B; 1.5 min, 85% B; 5.5 min, 35% B; 10 min, 35% B; 10.5 min, 35% B; 10.6 min, 10% B; 12.5 min, 10% B; 13.5 min, 85% B; and 20 min, 85% B. The flow rate was 0.15 ml/min from 0 to 5.5 min, 0.17 ml/min from 6.9 to 10.5 min, 0.3 ml/min from 10.6 to 17.9 min, and 0.15 ml/min from 18 to 20 min. All solvents are LC-MS grade and were purchased from Fisher.

### Mass spectrometry

The Q Exactive Plus mass spectrometer (Thermo Scientific) is equipped with a heated electrospray ionization probe, and the relevant parameters are listed: evaporation temperature, 120 °C; sheath gas, 30; auxiliary gas, 10; sweep gas, 3; spray voltage, 3.6 kV for positive mode and 2.5 kV for negative mode. Capillary temperature was set at 320 °C, and S lens was 55. A full scan range from 70 to 900 (m/z) was used. The resolution was set at 70,000. The maximum injection time was 200 ms. Automated gain control was targeted at 3 x 10^6^ ions.

### Peak extraction and data analysis

Raw data collected from LC-Q Exactive Plus MS was processed on Sieve 2.0 (Thermo Scientific). Peak alignment and detection were performed according to the protocol described by Thermo Scientific. For a targeted metabolite analysis, the method “peak alignment and frame extraction” was applied. An input file of theoretical m/z and detected retention time was used for targeted metabolite analysis, and the m/z width was set to 5 ppm. An output file including detected m/z and relative intensity in different samples was obtained after data processing. If the lowest integrated mass spectrometer signal (MS intensity) was less than 1,000 and the highest signal was less than 10,000, then this metabolite was considered below the detection limit and excluded for further data analysis. If the lowest signal was less than 1,000, but the highest signal was more than 10,000, then a value of 1,000 was imputed for the lowest signals. For isotope tracing experiments, the mass isotopomer distributions were calculated and normalized by comparing the ratio of labeled to unlabeled metabolites in each sample.

### Fly stocks and maintenance

*w^1118^* stocks were kindly provided by Dr. Don Fox. Flies were maintained on Nutri-fly food (Genesee Scientific 66-112) at room temperature.

### Fly Survival

Newly emerged adult male and virgin female flies were collected and sorted under CO2 anesthesia for three days prior to the start of survival experiments. On day 0, flies were again subjected to CO2 anesthesia to be randomized to their diet group and were put onto food containing their assigned diet (male and female flies were kept in separate vials with 30-40 flies/vial at the start of the experiment). During the experiment, flies were moved to a new vial of food every 3-4 days. Vials were visually inspected for dead flies and deaths were recorded daily each week from Monday-Friday. Flies were reared on benchtops at room temperature, with each different diet group subjected to the same conditions.

### *Drosophila* Chemically Defined Diets

Drosophila diet formulations have been previously published^51^ and were derived from previous recipes^52,53^ with the following modifications: (1) the type of Agar (Micropropagation Agar-Type II; Caisson Laboratories #A037), (2) the final percentage of Agar (1%), 3) the amount of sucrose (25 g), 4) the amino acids that were added to stock solutions before or after autoclaving^54^ whose order is described below and 5) the exclusion of inosine and uridine from the all diets except for the “cysteine and methionine free + nucleosides diet”, which contained 2.5 mM inosine, uridine, adenosine, guanosine, cytidine, and thymidine added into the “Other Nutrients” solution. The amino acid composition of the diet was based on the exome-matched (i.e. the concentrations used for a given amino acid correspond with the prevalence of exons for that amino acid in the Drosophila genome) Drosophila diet formulation developed in a previous study^53^ that was found to be optimal for growth and fecundity without compromising lifespan. The rationale for which amino acids were part of the autoclaving process was based on solubility considerations^54^.

The complete procedure, formula, and stock solutions for food production are as follows: (Note: the procedure below was used to create the “control diet”. The cysteine and methionine depleted diets were created the same way, but without the addition of cysteine and/or methionine. For these diets, instead of adding the cysteine or methionine stock solution, we added an equal volume of the solution the amino acid would be suspended in to match water content and pH.)

#### Procedure

1. Prepare “Part 2” (see below) Mixture and set aside;
2. Prepare “Part 1” (see below) Mixture, adding everything but agar (not everything will go into solution at this point);
3. Add agar to “Part 1” Mixture, stir using stir bar;
4. Autoclave “Part 1” Mixture for 15 minutes;
5. Remove “Part 1” Mixture from autoclave, then combine with “Part 2” Mixture and stir;
6. Quickly pipette food into Drosophila vials (5-10 mL food/vial);
7. Allow food to solidify/cool for roughly an hour, then cover vials (either with cotton plugs or with plastic wrap) and store food at 4°C.

Food is good for about one month at 4°C (will shrink and pull away from sides of vials due to loss of water after this).

#### Note

After autoclaving, “Part 1” Mixture containing agar can start solidifying (both before and after the two mixtures are combined, but combining the two mixtures will cause food to cool down quite a bit and solidify faster). Quickly combine and pour food while autoclaved mixture is still hot to avoid this. Adding water to the autoclave tray and keeping the “Part 1” Mixture in this hot water until ready to combine and pour helps keep it hot and helps prevent premature solidification.

##### Formula

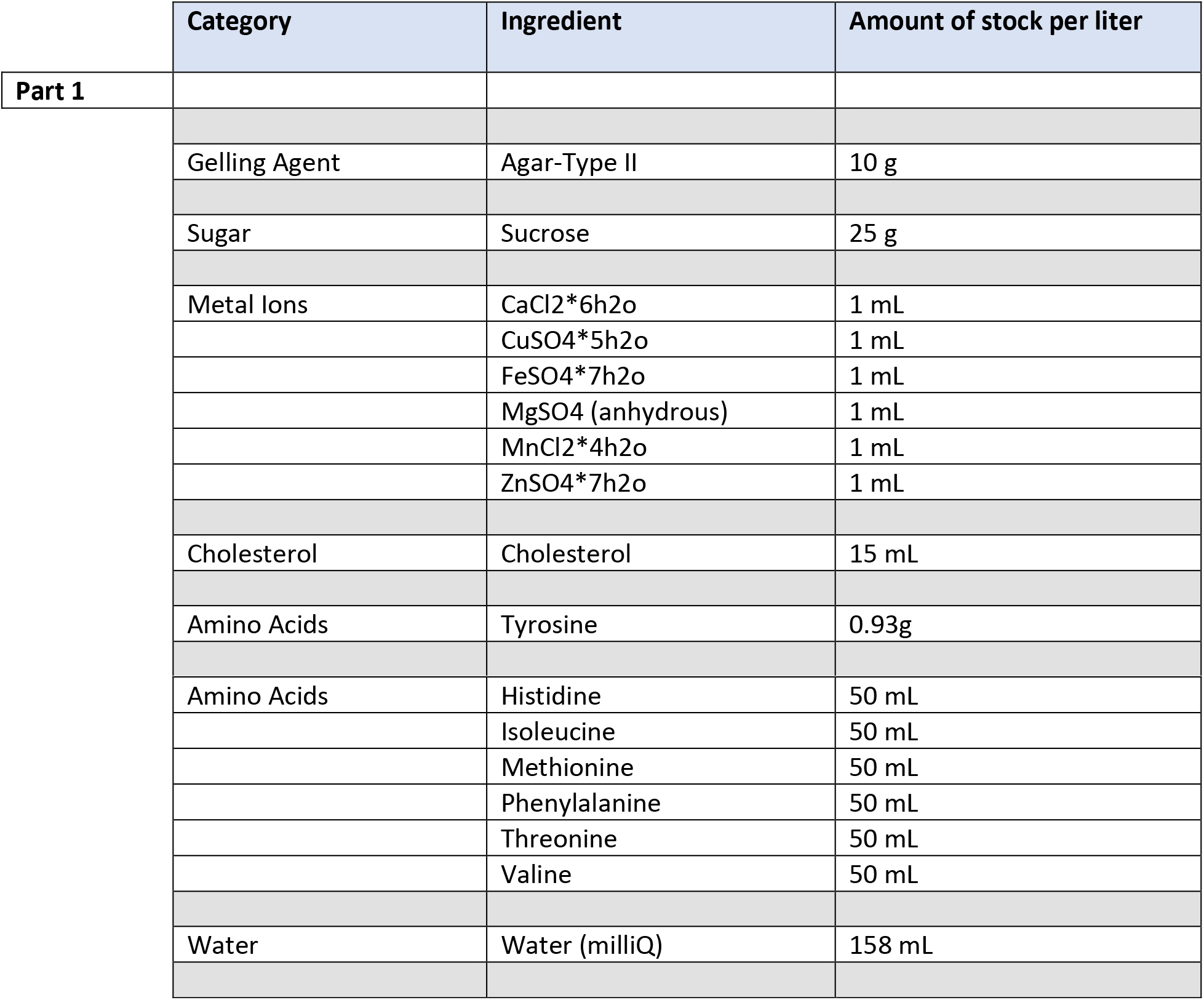

##### **AUTOCLAVE 15 minutes

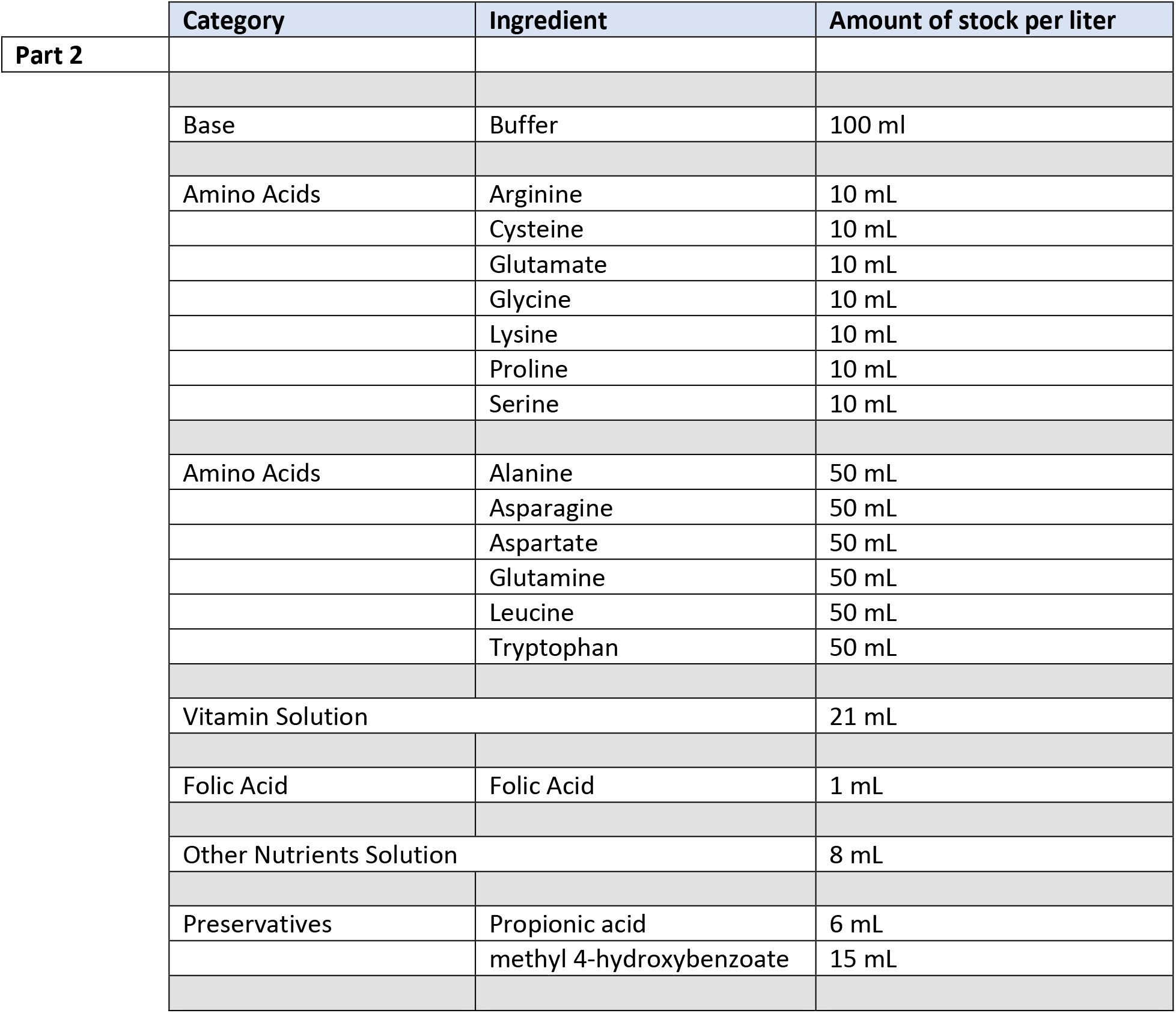

**\*\*AUTOCLAVE 15 minutes**
|  | Category | Ingredient | Amount of stock per liter |
| --- | --- | --- | --- |
| Part 2 |  |  |  |
|  | Base | Buffer | 100 ml |
|  | Amino Acids | Arginine | 10 mL |
|  |  | Cysteine | 10 mL |
|  |  | Glutamate | 10 mL |
|  |  | Glycine | 10 mL |
|  |  | Lysine | 10 mL |
|  |  | Proline | 10 mL |
|  |  | Serine | 10 mL |
|  | Amino Acids | Alanine | 50 mL |
|  |  | Asparagine | 50 mL |
|  |  | Aspartate | 50 mL |
|  |  | Glutamine | 50 mL |
|  |  | Leucine | 50 mL |
|  |  | Tryptophan | 50 mL |
|  | Vitamin Solution |  | 21 mL |
|  | Folic Acid | Folic Acid | 1 mL |
|  | Other Nutrients Solution |  | 8 mL |
|  | Preservatives | Propionic acid | 6 mL |
|  |  | methyl 4-hydroxybenzoate | 15 mL |

##### Stock Solutions

| Amino Acids | Catalog Number | g/50mL | Suspend in: |
| --- | --- | --- | --- |
| L-Alanine | Sigma, A7469 | 1.10 | H2O |
| L-Asparagine | Amresco, 94341 | 1.03 | H2O |
| L-Aspartic Acid | Alfa Aesar, A13520 | 1.17 | 0.5N NaOH |
| L-Glutamine | Amresco, 0374 | 1.12 | H2O |
| L-Histidine | Amresco, 1B1164 | 0.65 | H2O |
| L-Isoleucine | Amresco, E803 | 1.12 | H2O |
| L-Leucine | Sigma, L8912 | 2.03 | 0.2N HCl |
| L-Methionine | Amresco, E801 | 0.60 | H2O |
| L-Phenylalanine | Sigma, P5482 | 1.01 | H2O |
| L-Threonine | Sigma, T8441 | 1.11 | H2O |
| L-Tryptophan | Amresco, E800 | 0.32 | H2O |
| L-Valine | Amresco, 1B1102 | 1.20 | H2O |
| L-Arginine HCl | Amresco, 0877 | 8.16 | H2O |
| L-Cysteine | Sigma, 30089 | 1.71 | 1N HCl |
| L-Glutamic acid | Alfa Aesar, A12919 | 7.59 | H2O |
| L-Glycine | Alfa Aesar, A13816 | 3.84 | H2O |
| L-Lysine HCl | Amresco, 0437 | 6.83 | H2O |
| L-Proline | Sigma, P5607 | 4.89 | H2O |
| L-Serine | Sigma, S4311 | 6.89 | H2O |
| L-Tyrosine | Sigma, T8566 | **add Tyr powder |  |
| **Vitamin solution** | **Catalog Number** | **g/50mL** | **Suspend in:** |
| Biotin | Sigma, B4501 | 0.001 | H2O |
| Ca pantothenate | Sigma, 21210 | 0.039 | H2O |
| Nicotinic acid | Sigma, N4126 | 0.030 | H2O |
| Pyridoxine HCl | Sigma, P9755 | 0.006 | H2O |
| Riboflavin | Sigma, R4500 | 0.003 | H2O |
| Thiamine (aneurin) | Sigma, T4625 | 0.005 | H2O |
| **Folic acid solution** | **Catalog Number** | **g/50mL** | **Suspend in:** |
| Folic acid | Sigma F8758 | 0.0250 | 0.004N NaOH |
| **Other Nutrients solution** | **Catalog Number** | **g/50mL** | **Suspend in:** |
| Choline chloride | MP Biomedicals, 194639 | 0.3125 | H2O |
| Myo-Inositol | Sigma, I7508 | 0.0315 | H2O |
| **Methyl 4-hydroxybenzoate solution** | **Catalog Number** | **g/50mL** | **Suspend in:** |
| Methyl 4-hydroxybenzoate | Sigma, H3647 | 5.0 | 95% EtOH |
| **Buffer** | **Catalog Number** | **50mL stock** |  |
| Glacial Acetic Acid | Millipore, AX0074 | 1.5 mL |  |
| KH2PO4 | JT Baker, 3246 | 1.5 g |  |
| NaHCO3 | Sigma, S8875 | 0.5 g |  |
| Water |  | Up to 50mL |  |
| **Metal Ions** | **Catalog Number** | **g/50mL** | **Suspend in:** |
| CaCl2*6h2o | Sigma, 21108 | 12.5 | H2O |
| CuSO4*5h2o | Sigma, C7631 | 0.125 | H2O |
| FeSO4*7h2o | Sigma, F7002 | 1.25 | H2O (store -20C) |
| MgSO<sub>4</sub> (anhydrous) | Sigma, M7506 | 12.5 | H<sub>2</sub>O |
| MnCl<sub>2</sub>*4h<sub>2</sub>o | Sigma, M3634 | 0.05 | H<sub>2</sub>O |
| ZnSO<sub>4</sub>*7h<sub>2</sub>o | Sigma, Z0251 | 1.25 | H<sub>2</sub>O |
| **Cholesterol solution** | **Catalog Number** | **g/50mL** | **Suspend in:** |
| Cholesterol | Sigma, C8253 | 1 | EtOH |

#### Catalog Numbers for other reagents

Sucrose: Sigma, S7903

Agar: Caisson, A037

Propionic acid: Sigma, P5561

Inosine: Sigma, I4125

Uridine: Sigma,U3003

Adenosine: Sigma, A4036

Guanosine: Sigma, G6264

Cytidine: Sigma, C4654

Thymidine: Sigma, T1895

Stocks can be stored at 4°C for several months unless otherwise specified.

### Statistics

Details on individual statistical tests are described in figure legends. Unless stated otherwise, bar graphs plot the mean and error bars show standard deviation. Individual dots represent one biological replicate.

## Acknowledgments

JWL appreciates funding support from the American Cancer Society (129832-RSG-16-214-01-TBE) and the National Institutes of Health (R01CA193256). MAR was supported by a postdoctoral fellowship from the American Cancer Society (131615-PF-17-210-01-TBE). AEA was supported by a predoctoral fellowship from the National Cancer Institute (F31CA232658) and the National Institute of Health’s (NIH) Pharmacological Science Training Program grant (5T32GM007105). JWL advises Restoration Foodworks, Petri Biologicals, Cornerstone Pharmaceuticals, and Nanocare Technologies. The authors declare no competing interests.

## Author Contributions

J.W.L. and A.E.A conceived the study, designed experiments, wrote and edited the manuscript, performed project administration, validation, and supervision. A.E.A., Y.S., and F.W. performed experiments and analyzed the data. M.A.R. adapted *Drosophila* chemically defined diets from previous formulations. A.E.A performed formal analysis and visualization. J.W.L., A.E.A., and M.A.R. acquired funding.

## Data and Materials Availability

All metabolomics datasets are available on GITHUB via https://github.com/LocasaleLab/Allen-et-al-2022.git. All other data needed to evaluate the conclusions in the paper are present in the paper and/or the Supplementary Materials. Additional data related to this paper may be requested from the authors.

## Supplemental Figures

**Supplementary Figure 1.**
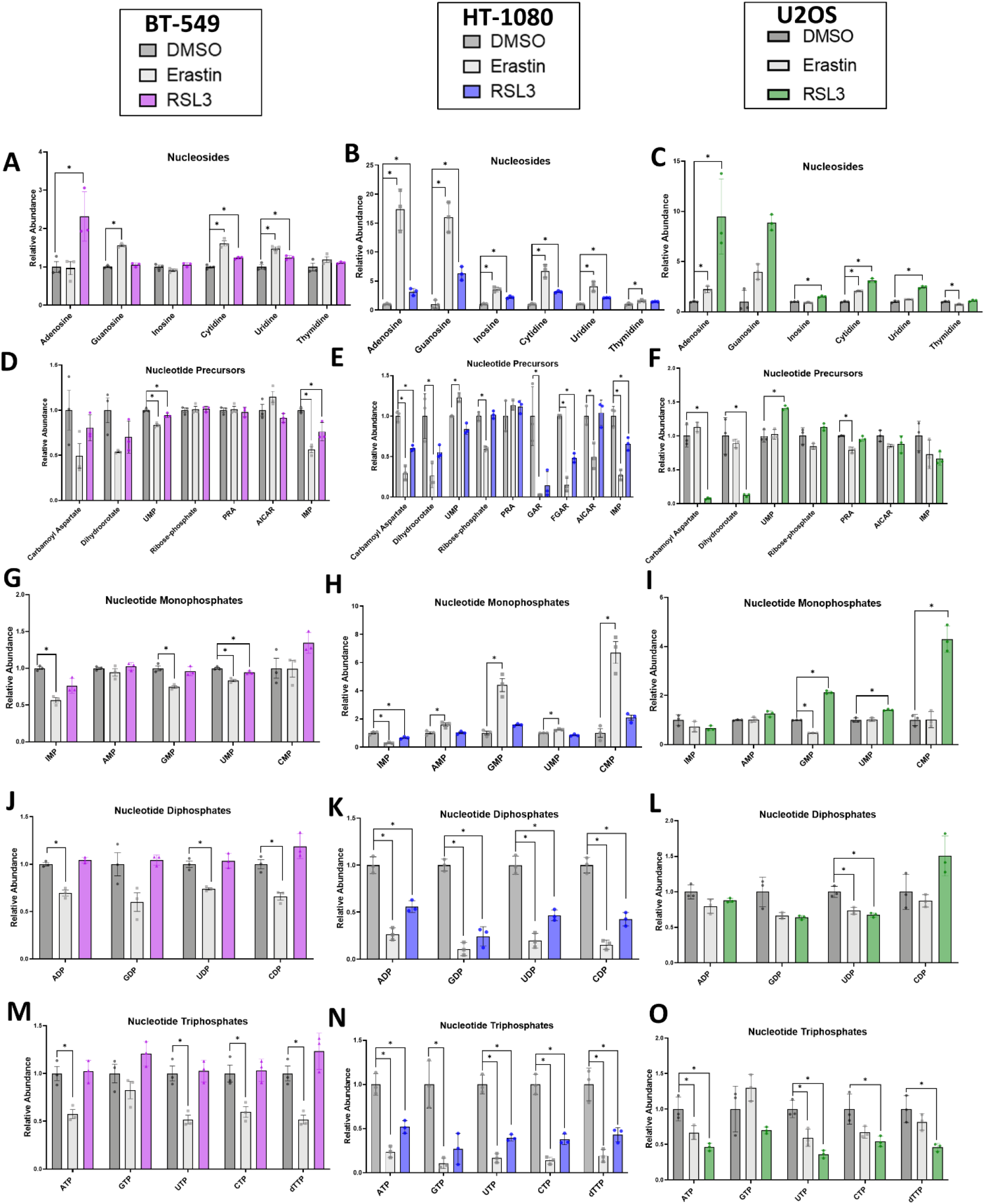
A, D, G, J, M) Nucleotide-related metabolites from BT-549 cells treated with 3.5 μM erastin or 75 nM RSL3. B, E, H, K, N) Nucleotide-related metabolites from HT-1080 cells treated with 2.5 μM erastin or 75 nM RSL3. C F, I, L, O) Nucleotide-related metabolites from U2OS cells treated with 5 μM erastin or 1 μM RSL3. For Figs. A-O, a two-way repeated measures ANOVA was performed on log-transformed values and followed by an uncorrected Fisher’s LSD test if interaction term or main effect of treatment p<0.05. * indicates p<0.05 when comparing DMSO to Erastin or RSL3 treatment in Fisher’s LSD test. Abbreviations: N-carbamoyl-L-aspartate (Carbamoyl aspartate), 5-phosphoribosylamine (PRA), Glycineamide ribonucleotide (GAR), Formylglycinamide ribonucleotide (FGAR), 5-amino-4-imidazolecarboxamide (AICAR)

**Supplementary Figure 2.**
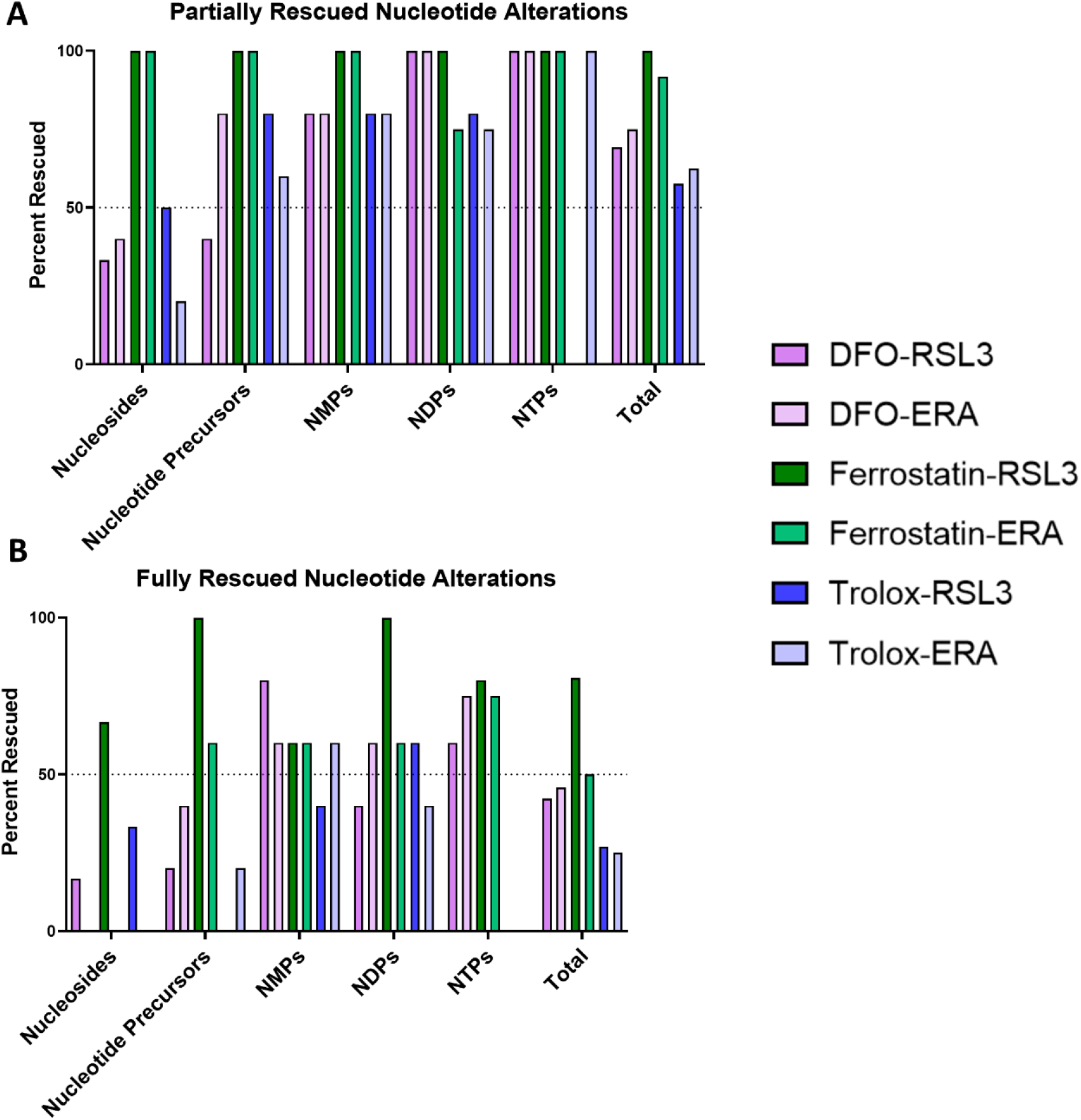
A) Percentage of nucleotide-related metabolites that were partially rescued by DFO, ferrostatin, or Trolox treatment in RSL3 or erastin (ERA)-treated HT-1080 cells. A partial rescue was defined as the rescue condition metabolite level being significantly higher or lower than in the non-rescue drug-treated condition (higher if erastin or RSL3 caused a significant depletion in the metabolite, and lower if they caused a significant accumulate of the metabolite). Significance was determined by a two-way repeated measures ANOVA performed on log-transformed values and followed by an uncorrected Fisher’s LSD test comparing drug-treated to control or drug-treated + rescue agent. B) Percentage of nucleotide-related metabolites that were fully rescued by DFO, ferrostatin, or Trolox treatment in RSL3 or erastin-treated HT-1080 cells. Metabolites were classified as “fully rescued” if they met both of the following conditions: 1) they were partially rescued defined by the conditions in A) and 2) the level of the metabolite in the rescue condition was not significantly different than the level of the metabolite in the control condition. Significance was determined by a two-way repeated measures ANOVA performed on log-transformed values and followed by an uncorrected Fisher’s LSD test comparing control to drug-treated + rescue agent.

**Supplementary Figure 3.**
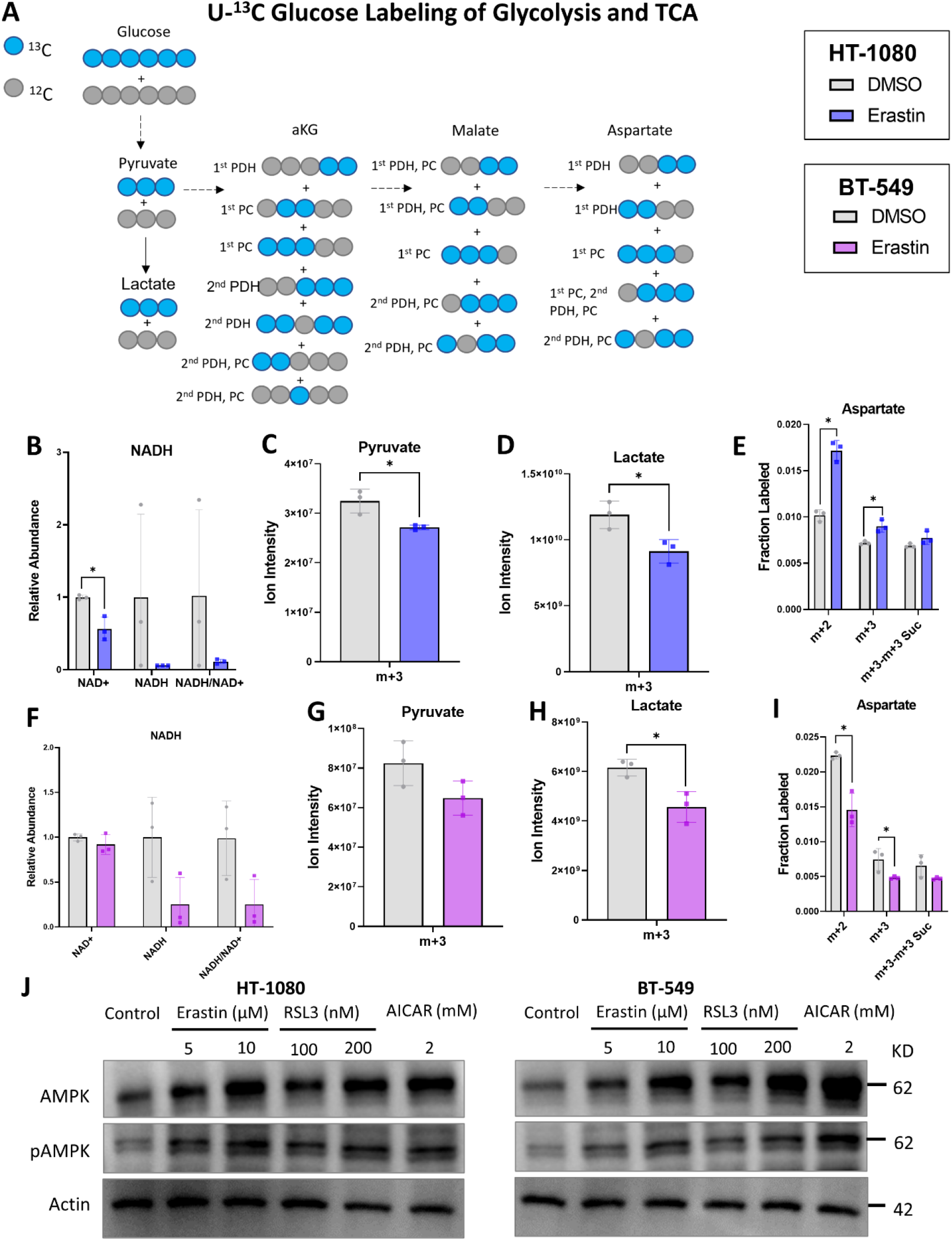
A) Schematic showing labeling of relevant glycolytic and TCA intermediates resulting from U-^13^C Glucose tracing. PDH means the TCA intermediate’s labeling pattern comes from pyruvate entering the TCA through pyruvate dehydrogenase and PC means the TCA intermediate’s labeling pattern comes from pyruvate entering the TCA though pyruvate carboxylase. 1^st^ means labeling from the 1^st^ turn of the TCA, 2^nd^ means labeling from the 2^nd^ turn of the TCA. B, F) NAD+ and NADH levels in HT-1080 cells treated with 2.5 μM erastin and BT-549 cells treated with 3.5 μM erastin. C, D, G, H) M+3 pool size for pyruvate and lactate from U-^13^C Glucose tracing. E, I) Aspartate fractional labeling from U-^13^C Glucose tracing. m+3-m+3 Suc represents the m+3 fraction of aspartate minus the m+3 fraction of succinate, which represents glucose entry into the TCA via pyruvate carboxylase. J) Western blot analysis of AMPK, pAMPK (Thr172), and actin protein expression levels after cells were treated for 8 hours. For Figs. B and F a two-way repeated measures ANOVA was performed on log-transformed values and followed by an uncorrected Fisher’s LSD test if interaction term or main effect of treatment p<0.05. * indicates p<0.05 when comparing DMSO to Erastin treatment in Fisher’s LSD test and no stars indicates p>0.05. For C-E and G-I an unpaired T-test was performed to compare DMSO to erastin treatment. If more than one isotopomer is shown on the graph, a separate uncorrected T-test was performed for each. * indicates p<0.05 for this test, no stars shown indicates p>0.05.

**Supplementary Figure 4.**
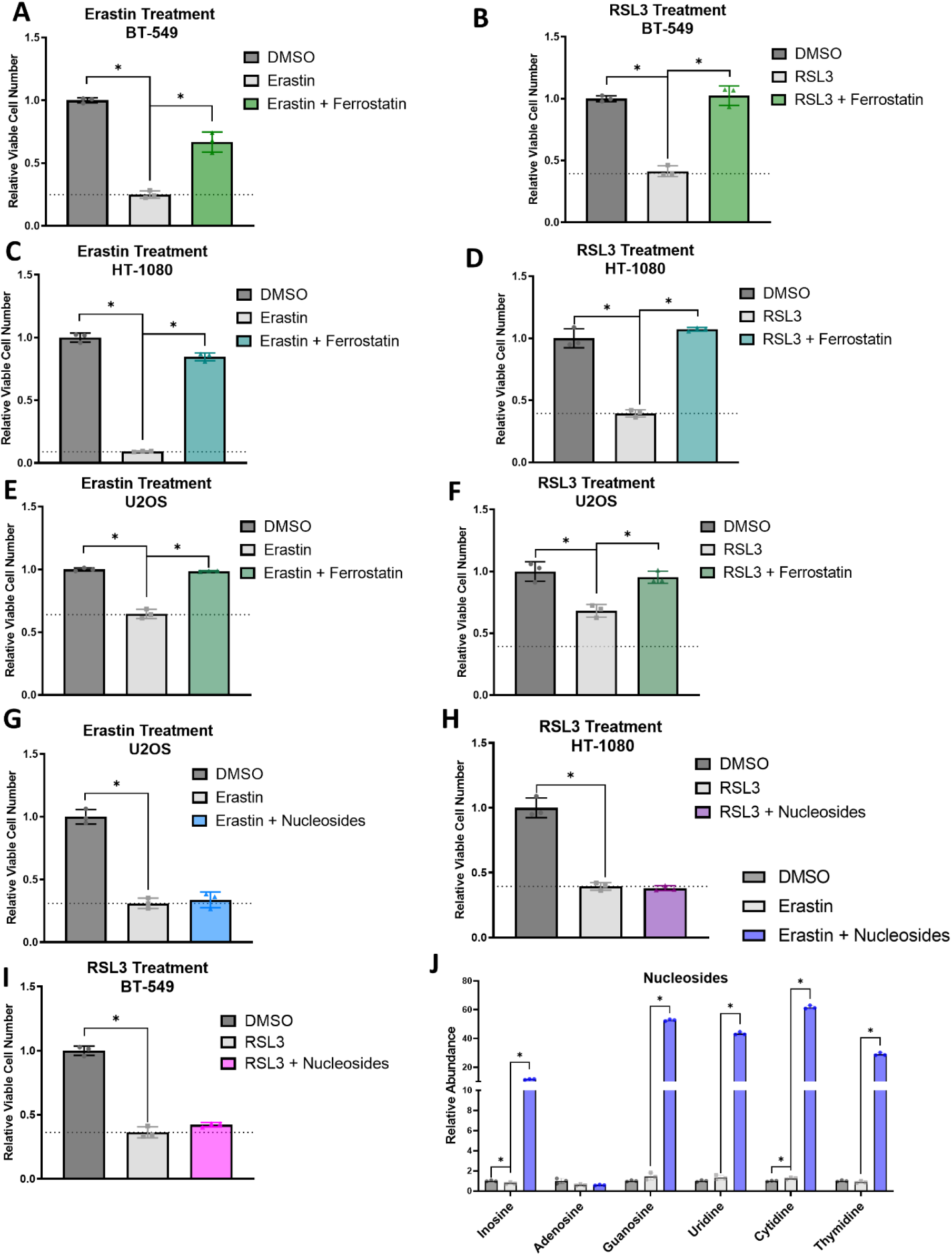
Relative viable cell number in: A) BT-549 cells treated +/- 3.5 μM erastin and 0.5 μM ferrostatin for 48 hours. B) BT-549 cells treated +/- 75 nM RSL3 and 0.5 μM ferrostatin for 48 hours. C) HT-1080 cells treated +/- 5 μM erastin and 0.5 μM ferrostatin for 48 hours. D) HT-1080 cells treated +/- 75 nM RSL3 and 0.5 uM Ferrostatin for 48 hours.. E) U2OS cells treated +/- 5 μM erastin and 0.5 μM ferrostatin for 48 hours. F) U2OS cells treated +/- 1 μM RSL3 and 0.5 μM ferrostatin for 48 hours. G) U2OS cells treated +/- 5 μM erastin and 1x nucleosides for 48 hours. H) HT-1080 cells treated +/- 75 nM RSL3 and 1x nucleosides for 48 hours I) BT-549 cells treated +/- 75 nM RSL3 and 1x nucleosides for 48 hours. J) Relative nucleoside levels from BT-549 cells treated with 3.5 μM erastin +/- nucleosides. For Figs. A-I a one-way ANOVA was performed and followed by an uncorrected Fisher’s LSD test when p<0.05 for ANOVA. * indicates p<0.05 in Fisher’s LSD test. For Fig. J, a two-way repeated measures ANOVA was performed on log-transformed values and followed by an uncorrected Fisher’s LSD test if interaction term or main effect of treatment p<0.05. * indicates p<0.05 when comparing Erastin to DMSO or to Erastin + Nucleosides. Bar is plotted at mean and error bars show standard deviation. Each dot represents one biological replicate.

**Supplementary Figure 5.**
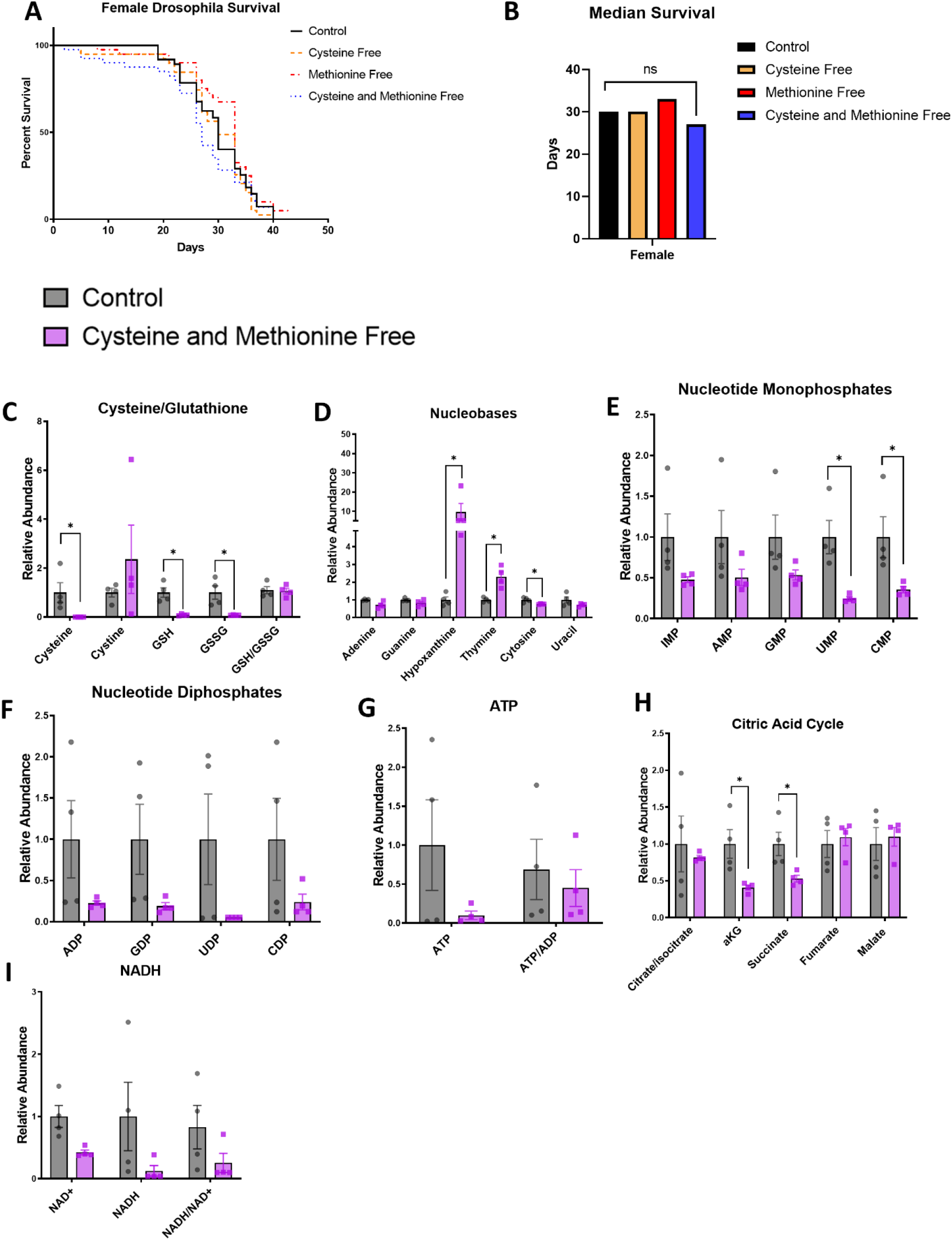
A) Virgin female *w1118 drosophila* survival when fed chemically defined diets with or without cysteine and methionine. B) Median survival of female flies from survival curves shown in Fig. S4A. C-I) The relative abundance of selected metabolites from male *drosophila* heads fed the indicated diets for 3 weeks. Two heads were pooled together, and each dot represents a pooled sample. For Fig. B, multiple log-rank Mantel-Cox tests were performed to compare female fly survival differences on control versus cysteine/methionine free diets. * indicates p<0.05 for these tests, while no star indicates p>0.05. For Figs. C-I, a two-way repeated measures ANOVA was performed on log-transformed values and followed by an uncorrected Fisher’s LSD test if interaction term <0.05. * indicates p<0.05 for Fisher’s LSD test. Bars are plotted at mean and error bars show standard deviation.

